# Heterogeneous T cell motility behaviors emerge from a coupling between speed and turning *in vivo*

**DOI:** 10.1101/785964

**Authors:** Elizabeth R. Jerison, Stephen R. Quake

## Abstract

T cells *in vivo* migrate primarily via undirected random walks, but it remains unresolved how these random walks generate an efficient search. Here, we use light sheet microscopy of T cells in the larval zebrafish as a model system to study motility across large populations of cells over hours in their native context. We show that cell-to-cell variability is amplified by a correlation between speed and directional persistence, generating a characteristic cell behavioral manifold that is preserved under a perturbation to cell speeds, and seen in Mouse T cells and *Dictyostelium*. These results suggest that there is a single variable underlying ameboid cell motility that jointly controls speed and turning. This coupling explains behavioral heterogeneity in diverse systems and allows cells to access a broad range of length scales.

## Introduction

Many immune cells migrate through tissue in search of antigen or pathogens. In some cases, such as during extravasation from blood vessels and homing to target organs, this migration is guided by chemokine gradients (***Witt et al., 2005***; ***Okada et al., 2005***; ***Germain et al., 2012***; ***Sarris and Sixt, 2015***). However, for naive T cells within T cell zones, *in situ* imaging studies have found that unguided random walk processes dominate ((***Miller et al., 2002***, ***2003***; ***Preston et al., 2006***; ***Cahalan and Parker, 2008***; ***Beltman et al., 2007***; ***Banigan et al., 2015***; ***Harris et al., 2012***; ***Worbs et al., 2007***; ***Textor et al., 2011***; ***Beauchemin et al., 2007***; ***Mrass et al., 2006***; ***Katakai et al., 2013***; ***Mrass et al., 2017***), reviewed in (***Mrass et al., 2010***; ***Krummel et al., 2016***)). This observation creates a conceptual challenge: T cells must dwell at scales of microns to make contact with antigen presenting cells (***Wülfing et al., 1997***; ***Krummel et al., 2000***; ***Beltman et al., 2009a***; ***Fricke et al., 2016***), yet migrate over scales of millimeters to find rare targets. A conventional diffusive random walk struggles to access these varied scales efficiently, since a walker that dwells near another cell for 1 minute would require several days to travel 1 mm. Several authors have suggested that T cells may have an intrinsic behavioral program that allows them to explore over different length scales (***Harris et al., 2012***; ***Krummel et al., 2016***; ***Mempel et al., 2004***). However, testing this hypothesis via *in situ* fluorescence microscopy raises inherent technical challenges: to observe a single cell accessing a broad range of spatial scales, it is necessary to have micron scale resolution over fields of view of millimeters, with low enough photodamage to observe the same cells at high spatiotemporal resolution over long periods. For example, one intriguing proposal is that T cells perform Levy flight (***Harris et al., 2012***), an anomalous random walk characterized by a power-law distribution of step sizes. Such random walks have been described in detail in the physics and ecology literature (***Shlesinger et al., 1995***; ***Bartumeus et al., 2005***; ***Viswanathan et al., 2011***), and their scale-free behavior provides a natural way for foragers to accelerate searches in many contexts (***Bartumeus et al., 2002***). However, observation over short periods cannot distinguish between Levy flight and heterogeneity amongst individual walkers (***Petrovskii et al., 2011***), both of which can create a broad distribution of displacements. More generally, we would like to understand whether there is a statistically-consistent behavioral program carried out by these cells.

To address this question, we used selective plane illumination microscopy (***Pitrone et al., 2013***; ***Power and Huisken, 2017***) to observe the native population of T cells in the live larval zebrafish (Tg(*lck*:GFP, *nacre*^−/−^ (***Langenau et al., 2004***)), over millimeter fields of view and periods of a few hours. We observed a population of motile cells in the tail fin and larval fin fold (***Figure 1A***, ***Figure 1***-***video 1***). We used this model system to dissect variation in cell behavior in a simple tissue context. Rather than a single broad distribution of speeds sampled by all cells, as in Levy flight, we observed considerable heterogeneity in both speed and turning behavior across cells. This observation prompted us to analyze the distribution of cell behaviors in a space defined by speed and turning statistics. Surprisingly, cell behaviors fell on a one dimensional manifold in this space, characterized by a coupling between speed and directional persistence. Analysis of previously-published data in Mouse T cells (***Gérard et al., 2014***) and *Dictyostelium* (***Dang et al., 2013***) within this framework showed that their migration statistics fell along a similar manifold. Our results suggest that a coupling between speed and turning may be an intrinsic feature of ameboid cell migration, that explains apparent heterogeneity in migration behavior in diverse systems and generates exploration at many length scales. This framework also predicts global regulation in the actin remodeling machinery underlying ameboid migration, such that diverse perturbations modulate one underlying control variable.

**Figure 1.**
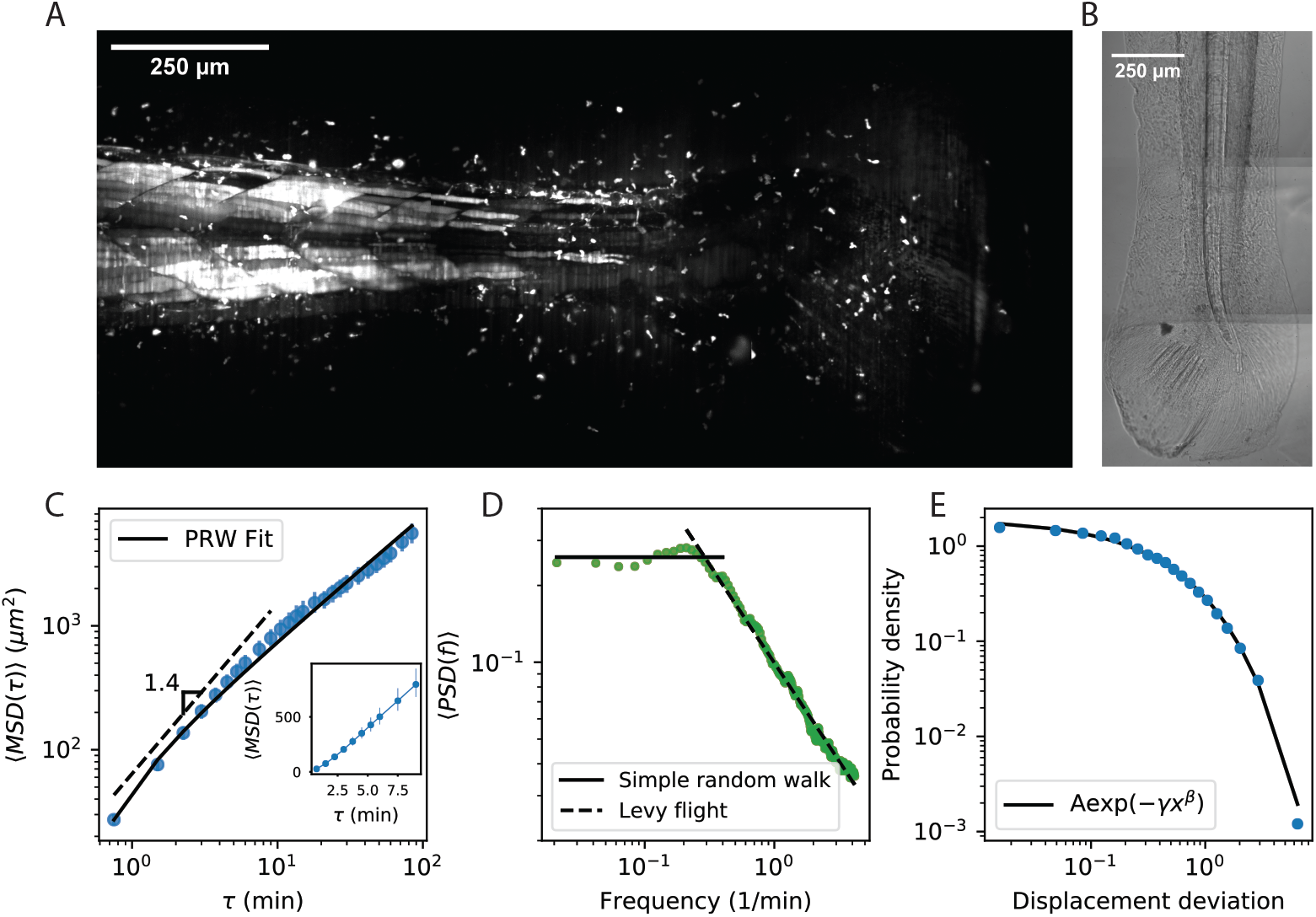
Cell motility behavior is inconsistent with Levy light. A. Maximum Z projection of a Tg(*lck*:GFP, *nacre*^−/−^) zebrafish at 12 dpf. This projection represents the first frame of a timecourse; see ***Figure 1***-***video 1***. B. Brightfield of the region of tissue shown in A. C. Mean squared displacement as a function of time lag. The cells migrate super-diffusively on scales of a few minutes. The MSD for a persistent random walk is fit to the data (Materials and Methods, Appendix 1). Error bars represent 95% confidence intervals on a bootstrap over n=335 trajectories containing all measured time intervals. (See also ***Figure Supplement 1***). Inset: linear scale for the first 10 minutes. D. The velocity power spectrum, averaged across all trajectories (n=634). A Levy (scale-free) process consistent with the short time behavior would result in a continuation of the high frequency slope (dashed line). Instead, we observe a timescale at a few minutes. E. Distribution of bout lengths within a trajectory (Materials and Methods), fit with a stretched exponential (n=36190 bouts). For all panels, trajectories were pooled from n=16 fish.

## Results

### Cell motility behavior is heterogeneous, inconsistent with Levy light

To investigate the statistical properties of T cell motility in our system, we measured cell trajectories within the tissue (Materials and Methods, ***Figure 1***-***video 1***, ***Figure 2***-***video 1***). We first evaluated evidence for Levy flight behavior, as opposed to persistent random walks (***Beauchemin et al., 2007***; ***Beltman et al., 2007***; ***Banigan et al., 2015***; ***Harris et al., 2012***), in our system. The distinction hinges on whether the statistics of individual trajectories are scale-free, so that super-diffusive behavior continues to long times; or if, alternatively, individual trajectories are diffusive at long times but there is heterogeneity across the population. To address this question, we performed a standard analysis of mean squared displacement as a function of time interval. Consistent with previous measurements (***Beauchemin et al., 2007***; ***Beltman et al., 2007***; ***Banigan et al., 2015***; ***Harris et al., 2012***), we observed a faster-than-linear increase in MSD at early times, indicating super-diffusive behavior, with a best-fit line in surprisingly good quantitative agreement with previous observations up through 10 minutes (***Harris et al., 2012***; ***Fricke et al., 2016***) (***Figure 1***C). However, we observed a transition at the scale of minutes, consistent with persistent random walks, and inconsistent with Levy flight (also note the straight line on a linear scale, ***Figure 1***C inset, characteristic of diffusive behavior). Note that while we have examined the subset of longer trajectories to measure the behavior through an additional order of magnitude in time, this result also holds when examining all trajectories through 15 minutes (***Figure 1***-***Figure Supplement 1***). To further test for an intermediate timescale, we computed the velocity-velocity power spectrum, using secant-approximated velocities along each trajectory (Materials and Methods). This quantity captures the timescale at which the velocities become decorrelated, if it exists; for a Levy-flight process the same negative slope is observed at all frequencies (***Viswanathan et al., 2005***), while a persistent random walk model passes towards zero slope at low frequencies (***Viswanathan et al., 2005***; ***Pedersen et al., 2016***). Consistent with the MSD analysis, we observe two regimes, with a clear timescale on the order of minutes (***Figure 1***D). Finally, we computed the distribution of lengths between direction changes (bout lengths) within a trajectory (Materials and Methods), scaled by the average bout length as suggested in (***Petrovskii et al., 2011***), and did not observe the characteristic Levy-flight power law (***Figure 1***E).

Since we did not find support for Levy flight in our system, we next evaluated evidence for cell-to-cell heterogeneity. From examples of velocity traces (***Figure 2***A,C-F,***Figure 2***-***video 1***), we observed substantial variation in speed between cells, that can persist over spans of a few hours. These trajectories are not atypical: overall, 88% of trajectories have distributions of secant-approximated speeds that are inconsistent with the speed distribution pooled on all trajectories (KS test, *p* < .01). Interestingly, we also found significant heterogeneity in cell turning behavior: 67% of cells had turn angle distributions inconsistent with the overall distribution (KS test, *p* < .01).

**Figure 2.**
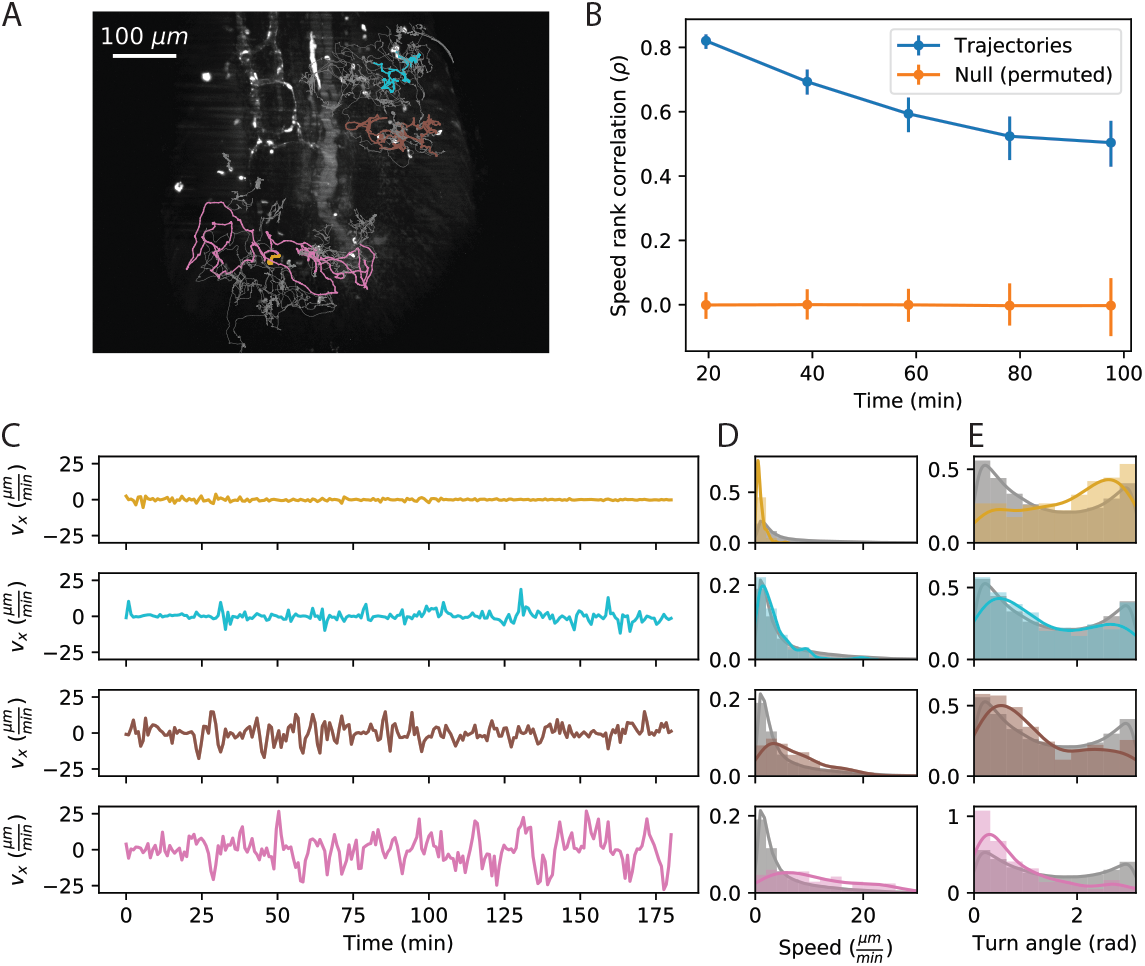
Cell speed and turning behavior are heterogeneous. A. Example of trajectories recorded over 3 hours at a 12 second interval (Tg(*lck*:GFP, *nacre*^−/−^) zebrafish; 10 dpf). Here we show a maximum Z projection of the 900th frame with trajectories overlaid; see ***video 1*** for the timecourse. Examples of four cell trajectories, with a range of characteristic speeds, are colored. B. Spearman rank correlation between trajectory speeds measured on non-overlapping 20 minute intervals, as a function of the time between the beginning of the intervals. Error bars represent 95% confidence intervals on a bootstrap over trajectories. The null model was constructed by permuting measured speeds across all the trajectories at each interval; error bars represent 95% confidence intervals over the permutations. (Calculations performed on the n=321 trajectories of at least 120 minutes in length.) C. Velocity traces for the four cells highlighted in A. D. Secant-approximated speed distributions for each cell from A, compared with the distribution over all cells (grey;n=98141 steps). E. Turn angle distributions for each cell from A, compared with the distribution over all cells (grey;n=96122 turn angles). Trajectories were pooled over n=16 fish.

To evaluate the rate of speed switching in our system, we measured the average speeds of individual trajectories on non-overlapping 20 minute intervals, and evaluated how the speed ranks change as a function of the time between intervals (***Figure 2***B). We found a high correlation between speeds on adjacent non-overlapping intervals, which decays slowly on the timescale of the measurement. Thus each cell samples a characteristic distribution of speeds that is stable over one to two hours. For the remainder of the analysis, we will consider the average speed to be a property of the trajectory; we return to consider the implications of speed switching in the discussion.

### Heterogeneous cell migration statistics fall on a behavioral manifold

The surprising degree of heterogeneity in random walk behavior amongst individual cells in this very simple tissue context led us to ask whether there are any underlying rules governing this variation. Are individual cells free to pick any turn and speed statistics, or are there constraints?

To investigate co-dependency between speed and turning behavior, we divided the cells into quintiles based on speed, which we refer to as speed classes. We observed strong variation in the distribution of turn angles amongst speed classes (***Figure 3***A): fast cells are most likely to turn shallowly, slow cells are most likely to turn around, and the distribution varies smoothly across the speed classes. This dependence could be driven by a local coupling between speed and turn angle: cells tend to go straighter whenever they go fast, which the faster cells do more often. Alternatively, it could be driven by an overall behavioral difference between fast and slow cells. To distinguish these possibilities, we measured the average turn angle as a function of the size of the steps surrounding it (***Figure 3***B). We found that both of these effects contribute: all cells go straighter during faster periods, but for a given step size, slow cells are more likely to turn sharply.

**Figure 3.**
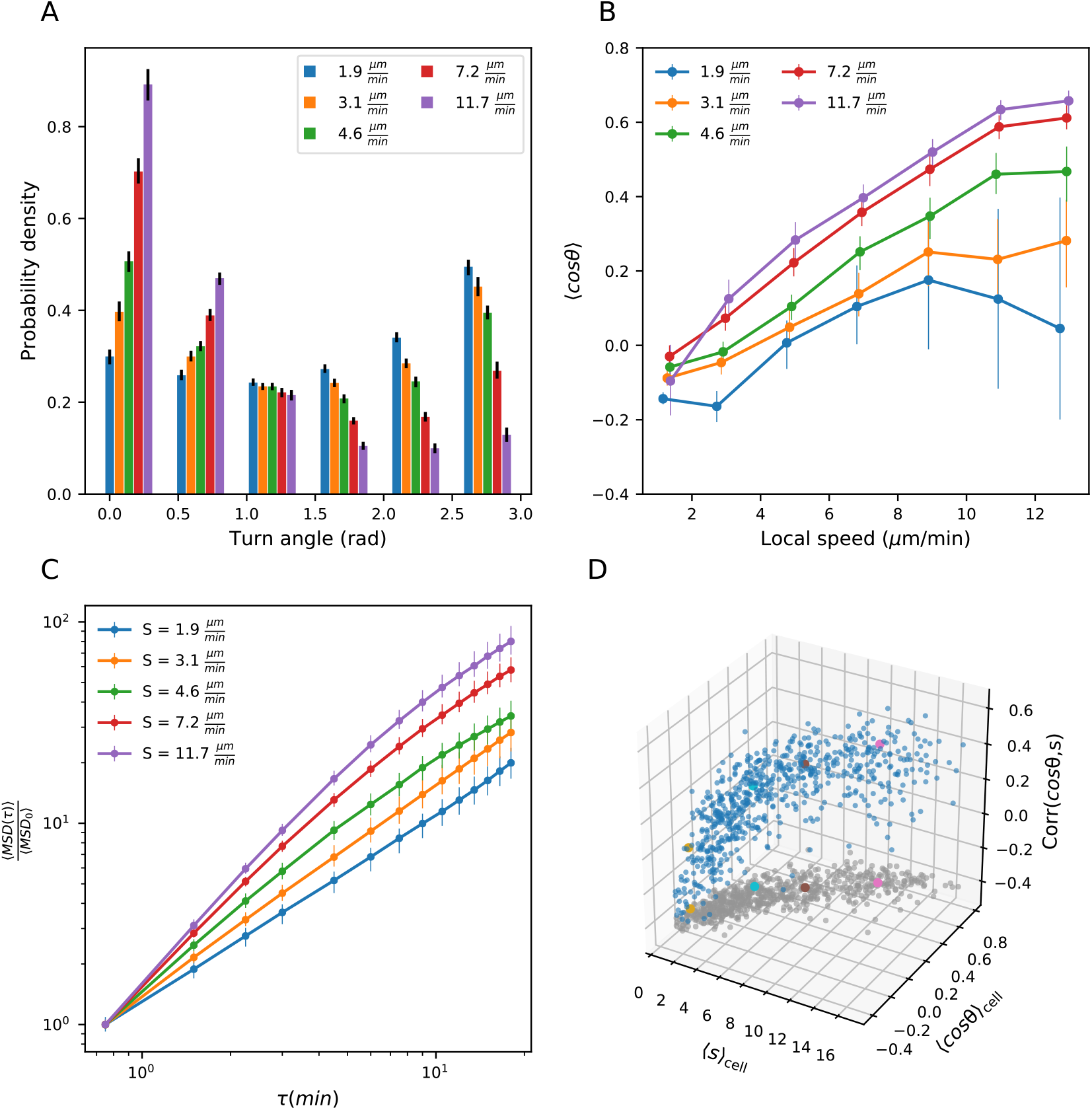
Heterogeneous cell migration statistics fall on a behavioral manifold. A. Distribution of turn angles amongst cells grouped by speed class. The distribution varies smoothly from faster cells, which tend to go straighter, to slower cells, which tend to turn around more often. Error bars represent 95% confidence intervals from a bootstrap over trajectories in each speed class. The legend reports the mean speed for trajectories in each class. B. Turning behavior conditioned on current cell speed. The average of the cosine of the turn angle as a function of the average length of the steps on either side. Cells are grouped into speed classes as in A. Error bars represent 95% confidence intervals from a bootstrap over trajectories in each speed class. C. Mean squared displacement by speed class. Due to the variation in turning behavior, the faster cells appear initially more superdiffusive. Error bars represent 95% confidence intervals from a bootstrap over trajectories in each speed class. All speed class calculations were performed on the n=569 trajectories that included all time intervals in the MSD analysis. D. Organization of cell behavior into a curve in a three dimensional behavioral space. Each point represents a trajectory, and we show the average speed, turn angle, and local speed-turn correlation. Grey: projection into the x-y plane. The trajectories shown in ***Figure 2*** are colored. Trajectories pooled over n=16 fish.

The relationship between speed and turning suggests that there may also be systematic differences in the scaling of the MSD at short times between cells. In particular, variation in speed alone amongst individuals would not change the shape of the MSD, which would collapse when appropriately scaled (Appendix 1). On the other hand, the systematically shallower turns of faster cells would be expected to boost the slope of their MSD at short times, an effect we observe in the data (***Figure 3***C).

The analysis at the level of speed classes suggested that there might be a single scalar variable, for which the cell’s average speed is a good proxy, that determines a number of higher-order statistics characterizing the cell’s migration behavior. To test this at the level of individual trajectories, we chose two summary statistics that capture the cell’s turning behavior: the average of the cosine of the turn angles along the trajectory, and the correlation between speeds and turn angles along the trajectory. The former is a summary of the overall distribution of turn angles for that cell, while the latter captures the degree of additional local coupling between speed and turn angle. Together with the cell’s speed, these two summary statistics form a three-dimensional behavioral space. We observed that the cell trajectories fall close to a curve in this space (***Figure 3***D). In particular, 73% of the variance in the average cosine can be explained by cell speed, with some residual variance due to the stochasticity of the process (7%) and other unknown effects (20%) (***Figure 3***-***Figure Supplement 1***). Thus T cell migration statistics can be organized into a one-dimensional behavioral manifold, characterized by a strong dependence between speed and turning behavior.

**Figure 2–video 1. Heterogeneity of T cell migration** Maximum Z projection of the tail of a Tg(*lck*:GFP, *nacre*^−/−^) at 10 dpf (GFP channel), with cell trajectories overlaid. A Z stack was recorded every 12 seconds for 3 hours (62 2 *μm* slices per stack). A maximum pixel value threshold of 1200 was used throughout the timecourse (no minimum pixel threshold was used). Four trajectories were chosen and highlighted in color; the remainder of the trajectories are plotted in grey. The movie was prepared using Python 3.6.0 (code available at: https://github.com/erjerison/TCellMigration).

### Model predicts wide variation in length scales of exploration across the population

Our observation of a behavioral manifold suggests that, despite the apparent heterogeneity in migration strategies, there may be a common program with a single underlying variable. In this view, a cell’s location on the manifold reflects its internal value of this control variable, which in turn dictates its random walk behavior. Given the results of our MSD analysis, to determine candidates for a single-parameter migration model, we started with the canonical persistent random walk (Ornstein-Uhlenbeck) process (***Uhlenbeck and Ornstein, 1930***)):

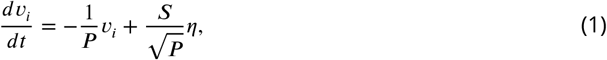

where v is the velocity, *η* is a white noise term, and *i* labels the velocity component. This model has two free parameters: the speed, S, and the persistence time, P, which is the average time before a cell turns. Our observations suggest that there may in fact only be one control parameter; in particular, because faster cells tend to make shallower turns, we expect P to increase with S. To determine the relationship between these two variables, we measured the persistence time, averaged along each trajectory, as a function of cell speed, and found a linear dependence (***Figure 4***A). This suggests the following simple model of cell motility:

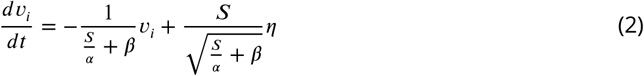

where *α* is a constant with units of acceleration, *β* is a constant with units of time, and both are constrained by the empirical relationship in ***Figure 4***A. We call this the speed-persistence coupling model (SPC).

**Figure 4.**
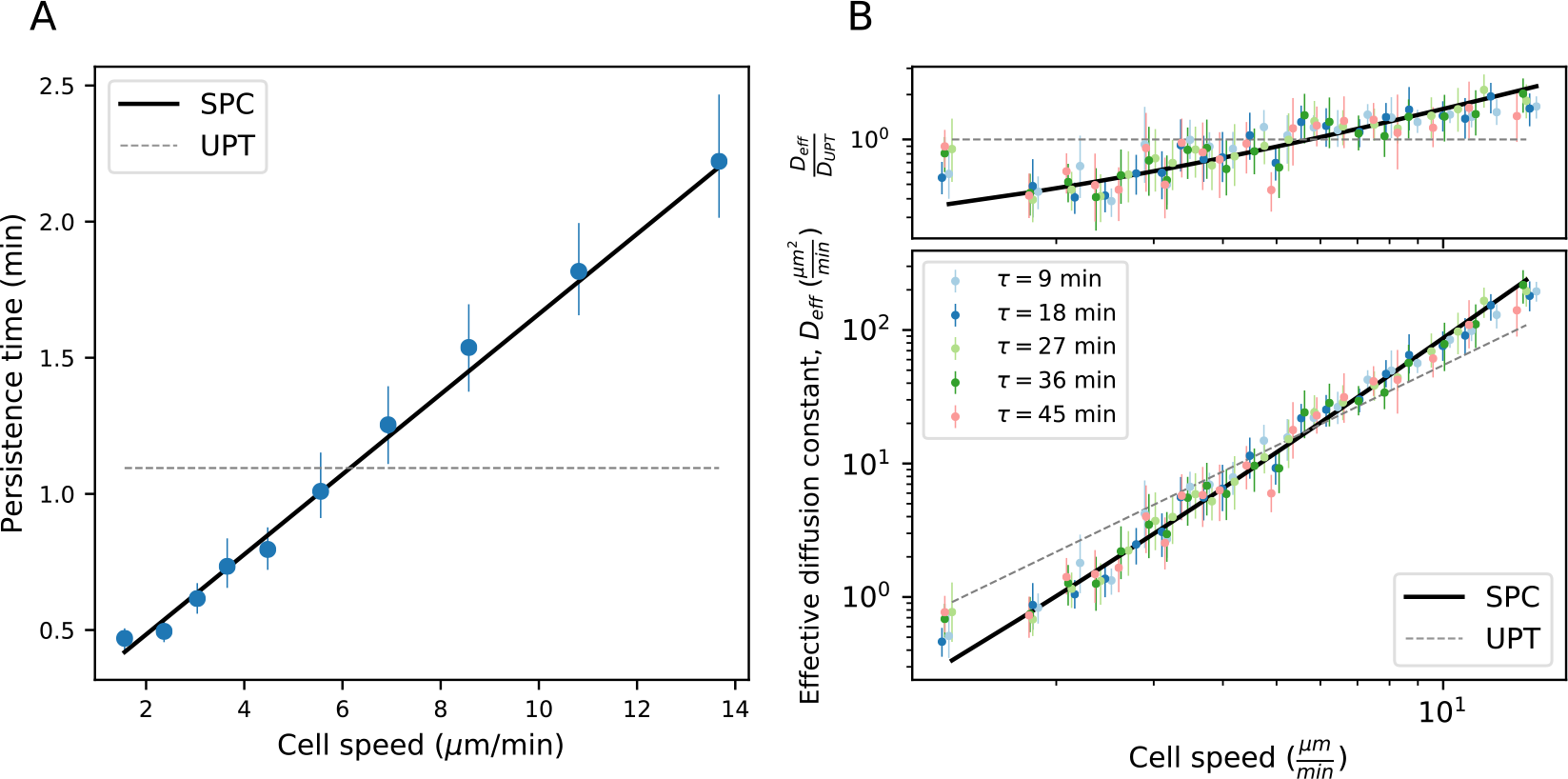
Model predicts wide variation in length scales of exploration across the population. A. Mean persistence time as a function of cell speed, measured along trajectories (n=710). Error bars represent 95% confidence intervals from a bootstrap over trajectories. UPT: Uniform persistence time; SPC: Speed-persistence coupling. B. Scaling of the effective diffusion constant with cell speed. Except for a constant offset, parameters are fixed based on the speed-persistence relationship in A. Error bars represent 95% confidence intervals on a bootstrap over trajectories. Numbers of trajectories in each time interval: n=704; n=654; n=607; n=558; n=523. Trajectories were pooled over n=16 fish.

As in other persistent random walk models, SPC walkers are diffusive at long times; the MSD scales linearly with time, and the ratio between these quantities defines an eff ective diff usion constant (Appendix 1):

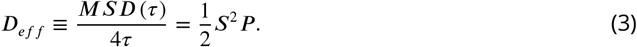

Due to the dependence of P on S, the SPC model predicts a strong scaling of the effective diffusion constant with cell speed. We tested this prediction at several time intervals *τ* and found good quantitative agreement between the model and the data (***Figure 4***B). In particular, the SPC model generates five-fold more variation in the effective diffusion constants across the cells than would be expected for a uniform persistence time model (UPT).

We note that the analyses in this and the previous section depend on measured cell speeds and turn angles, which are an imperfect proxy for the true instantaneous process (***Beltman et al., 2009b***). In particular, both noise in the cell locations and finite sampling intervals can introduce bias in the measured speeds, which could in principle generate spurious relationships between measured speed and turning behavior. We took two approaches to addressing the sensitivity of our conclusions to these issues. First, we addressed sensitivity to sampling rate by repeating the analyses above, subsampling timepoints by a factor of 2. This makes the turning behavior of the slowest two speed classes harder to distinguish, because they are rarely persistent over more than one timestep (***Figure 4***-***Figure Supplement 1***A,D), and introduces more noise in the local coupling (***Figure 4***-***Figure Supplement 1***B), but otherwise does not alter the structure of the correlations (***Figure 4***-***Figure Supplement 1***A-F). Second, we assessed the potential biases introduced by mislocation noise and finite sampling to the speed-persistence relationship in simulations (Appendix 1, Appendix 1-Figure 7). We found that mislocation noise can lead to spurious correlations between speed and persistence at the slow end of the speed spectrum, but cannot account for the consistent correlation we observe across speeds.

Finally, we note that the SPC Langevin model describes the effective diffusive behavior of the trajectories and their scaling at longer times, but may not capture all the details of the microscopic dynamics. In particular, the propensity of trajectories to turn backwards (peak at *θ* = *π* radians, ***Figure 3***A) is not captured by this model.

### Manifold is preserved under a drug perturbation to cell speeds

We next asked about the robustness of the observed behavioral manifold under a perturbation to cell speeds. To determine relevant pathways and candidates to perturb cell speed, we performed single-cell RNA sequencing on cells isolated from the tail of 15 dpf Tg(*lck*:GFP) zebrafish. To assess the fidelity of the marker, we sorted GFP+ cells from an unbiased FSC/BSC gate (Materials and Methods). We used standard dimensional reduction and clustering methods (Materials and Methods) to identify 351 putative T cells (***Figure 5***A-B). Unexpectedly, we also identified a population of epithelial cells that may mis-express *lck* at low levels (Materials and Methods, ***Figure 5***-***Figure Supplement 1***, ***Figure 5***-***Figure Supplement 2***).

**Figure 5.**
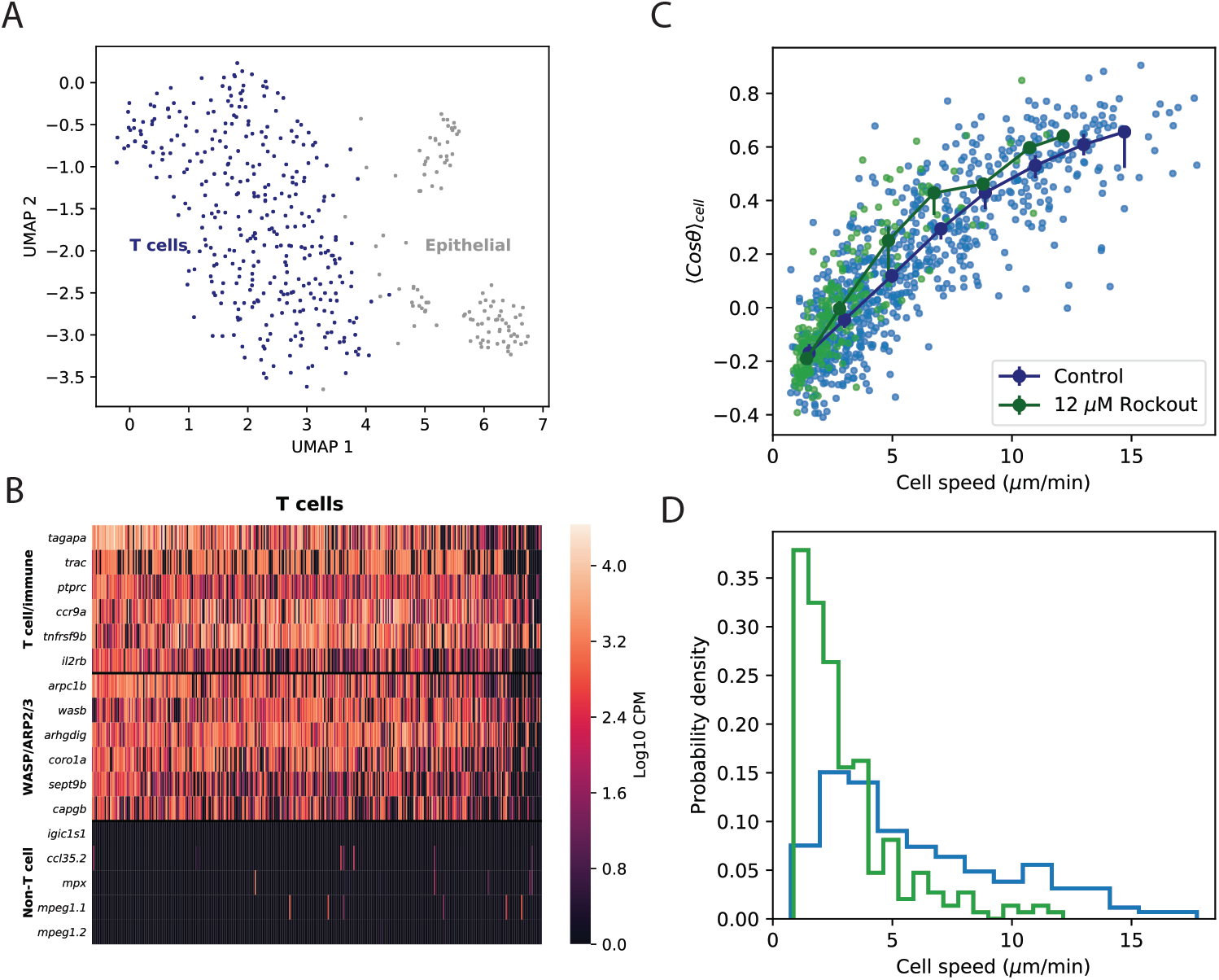
Manifold is preserved under a drug perturbation to cell speeds. A. Dimensional reduction via UMAP of scRNAseq gene expression profiles for cells sorted from 15 dpf Tg(*lck*:GFP) zebrafish. We called 351 putative T cells and 110 putative epithelial cells (Materials and Methods). B. Marker gene expression in the T cell cluster. Cells from the T cell cluster express a number of T cell and ubiquitous immune markers, including the T cell receptor light chain, trac, as well as genes involved in actin nucleation and remodeling. With the exception of trac, the genes shown in these first two categories were amongst the top 50 differentially expressed genes for the T cell cluster (Wilcoxon rank-sum test). Finally, we do not observe significant expression of immune markers associated with B cells, macrophages, or neutrophils. C. Correlation between the average cosine of the turn angles along the trajectory and cell speed, for cells in control and Rockout-treatment conditions. Data for all cells is shown as well as a binned average. Error bars represent 95% confidence intervals on the binned average on a bootstrap over cells. D. The distribution of speeds amongst control and Rockout-treated trajectories. The treatment lowers cell speeds but maintains the relationship between speed and persistence. Statistics based on trajectories pooled over n=16 control fish (n=712 trajectories) and n=6 Rockout treatment fish (n=236 trajectories). (See also ***Figure Supplement 3***.

We called marker genes for the putative T cells based on differential expression relative to the epithelial cells (Materials and Methods). These included a number of T cell and immune markers (***Schaum et al., 2018***; ***Moore et al., 2016***; ***Tang et al., 2017***) (***Figure 3***B). We note that we did not observe significant expression of markers associated with other motile non-T immune cells (B cells, macrophages, or neutrophils) (***Schaum et al., 2018***; ***Tang et al., 2017***), and we do not find support for NK cells (Materials and Methods, ***Figure 5***-***Figure Supplement 2***, ***Tang et al.*** (***2017***); ***Carmona et al. (2017***)). The T cell associated genes also included several canonically involved in actin nucleation and remodeling in the leukocyte cytoskeleton (***Vicente-Manzanares et al., 2002***; ***Takenawa and Suetsugu, 2007***) (WASP/ARP2/3 pathway; ***Figure 5***B, ***Figure 5***-***Figure Supplement 2***).

Based on these results, we chose the drug Rockout, a known Rho kinase inhibitor affecting this pathway (***Barros-Becker et al., 2017***), as a candidate for perturbing cell speed, and repeated the measurements and analysis of cell migration behavior in the presence of the drug (Materials and Methods). We found that the distribution of cell speeds shifted downwards, but we still observed a quantitatively similar positive relationship between speed and turning behavior (***Figure 5***C, D). This is consistent with a model where the perturbation primarily shifted an internal cell state variable that determines location along the behavioral manifold, which in turn dictates both speed and turning behavior, although we note that there may be an additional small shift towards shallower turns in the drug condition.

### Data from Mouse T cells and Dictyostelium also support speed-turn coupling

Finally, we analyzed published data from two other species, mouse T cells *in situ* (***Gérard et al., 2014***) and *Dictyostelium* (***Dang et al., 2013***), in this framework. While some of the analyses that depend on longer time traces and larger cell numbers are not possible with these datasets, we tested the relationship between average turn angle and cell speed, which drives many other differences in the dynamics. We found that this correlation held amongst the control cells in both studies (***Figure 6***). This suggests that, as for zebrafish T cells, there is heterogeneity in speed and turning behavior amongst the cells, and is consistent with a similar behavioral manifold. In the two published studies, genetic perturbations that knocked out or down one member of the actin remodeling machinery were used: a knockout of the non-canonical myosin Myo1g in one case, and a knockout of the Arp2/3 inhibitor Arpin in the other. In each case, the perturbation had a substantial effect on the distribution of cell speeds (***Figure 6***, C-D). However, in both cases, a quantitatively similar positive relationship between the speed and turning behavior amongst the perturbed cells was preserved.

This analysis, together with the effects of the small molecule perturbation used in this study, generate the prediction that there is global coupling amongst some components of the actin remodeling machinery, so that diverse perturbations can lead to similar internal states.

**Figure 6.**
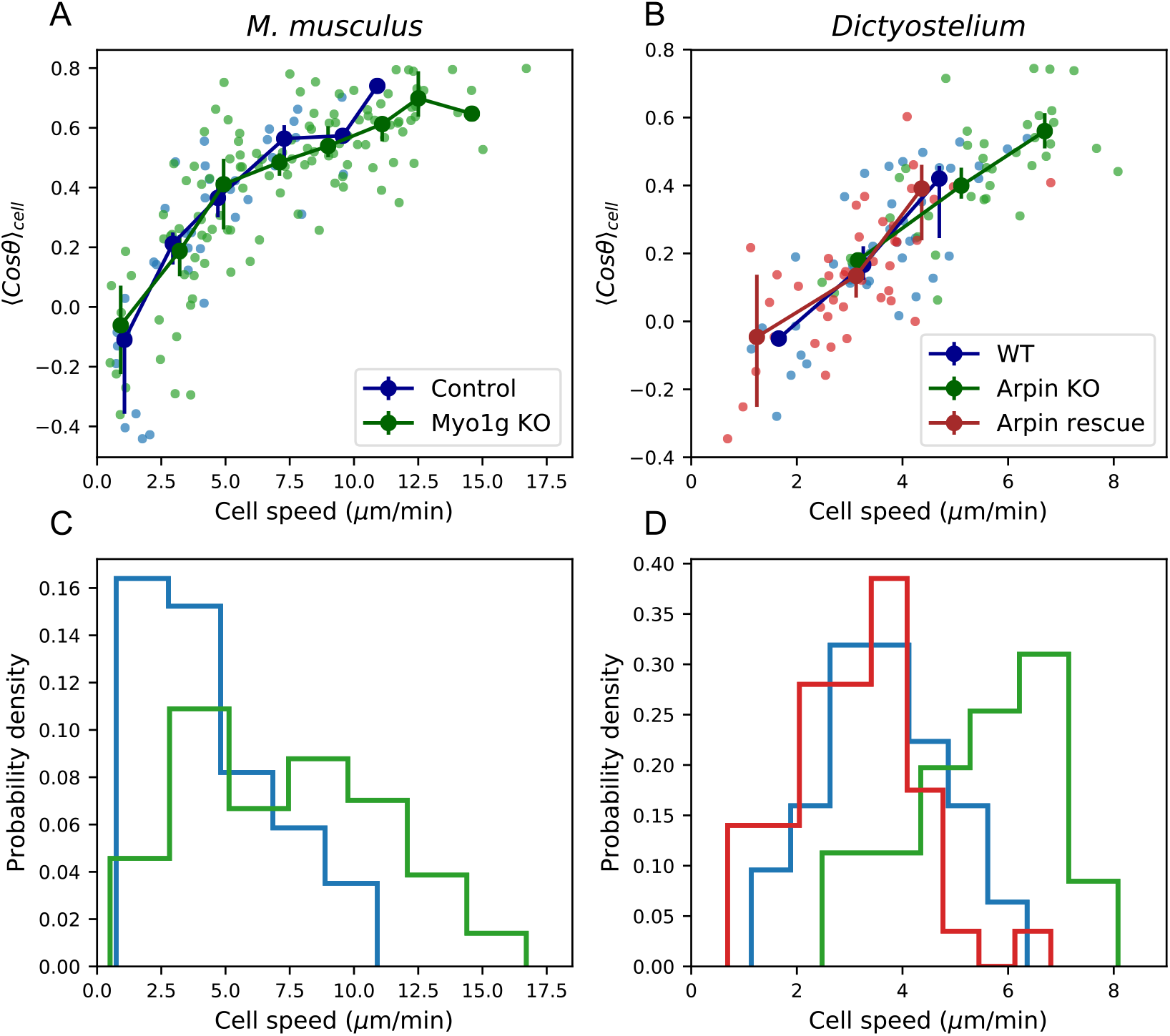
Data from Mouse T cells and *Dictyostelium* also support speed-turn coupling. A. As in 5B-C, for mouse T cells (data from (*Gérard et al., 2014*)). The perturbation is a genetic knockout of a non-canonical myosin motor, Myo1g. B. Same as in A, for *Dictyostelium* (data from (*Dang et al., 2013*)). The perturbations are a knockout and rescue of the Arp2/3 inhibitor Arpin. (control: n=42; Myo1g KO: n=123) C-D. Distributions of cell speeds for the control and treatment conditions shown in A-B. (WT: n=42; Arpin KO: n=38; Arpin rescue: n=42) In each case, the distribution of speeds shifts, but the cells tend to move along the speed-turn curve.

## Discussion

We have measured and analyzed the variability in cell motility amongst the T cells of the zebrafish tail, and used the framework generated by this analysis to examine Mouse T cells and *Dictoystelium*. We found that migration statistics from all three species fell on a similar behavioral manifold. We note that, in general, heterogeneity of motility behavior across a population could be caused by the tissue context rather than by cell-intrinsic factors. However, the effects of the drug perturbation, as well as the effects of the genetic perturbations from (***Gérard et al., 2014***) and (***Dang et al., 2013***), support a cell-intrinsic basis for the behavioral manifold we observe here. In particular, we performed trials in which the same regions of tissue were imaged and cells tracked before and after addition of the drug (***Figure 5***-***Figure Supplement 3***); the observed changes in migration statistics must then be caused by the drug’s effect on the cell’s internal state, not by the tissue context.

Our analysis of data from Mouse T cells and *Dictyostelium* suggests that speed-persistence coupling may be a general feature of ameboid cell migration. Most surprisingly, three apparently unrelated perturbations to the actin nucleation and remodeling machinery in the three different systems all had the effect of shifting cells along, rather than off, the manifold. This suggests that perturbations to different parts of the pathway may modulate a single underlying variable, which jointly controls speed and turn distributions. This generates the hypothesis of extensive regulated coupling within this pathway, such that there are relatively few true control variables.

The genetic perturbations used in previous studies made the cells faster and more persistent on average (***Gérard et al., 2014***; ***Dang et al., 2013***). Our results suggest that this connection may be general to the cells rather than specific to the perturbation. In particular, shifts in the average turning behavior have been used to argue that Arpin and Myo1g control cell steering. Our analysis suggests that increasing cell speed may in many cases increase straightness, and vice versa, so that the effect on cell steering may be indirect.

We have analyzed migration statistics in a framework where the average speed is a property of the trajectory. Our analysis of speed switching suggests that this is a reasonable assumption over about a 2 hour timespan (***Figure 2***B). We note that the moderate amount of speed switching that occurs within trajectories in our dataset may contribute to the unexplained portion of the variance in the speed-turn relationship (***Figure 3***-***Figure Supplement 1***). Our results suggest that cells maintain a location along the behavioral manifold, and hence an effective diffusion constant, over these timespans, so that there are metastable periods where cells explore at a particular length scale, before shifting to a different one. Because of the steep scaling of the effective diffusion constant with speed, the range of length scales accessible to the cells is large, with over 300-fold variation in effective diffusion constants in our data. Thus, while our observations are inconsistent with Levy flight, the SPC model generates other ways for the cells to explore across a broad span of scales.

Additionally, cells sampled from different parts of the manifold have different MSD slopes at early times (***Figure 3***C), so that this single-parameter migration model produces an apparent variety of migration strategies: if measured over short intervals, the cells appear to range from more to less superdiffusive.

Finally, we note that speed-turn coupling could enhance a search strategy where local chemokine cues cause cells to slow down (***Dustin et al., 1997***; ***Mempel et al., 2004***; ***Kawakami et al., 2005***; ***Castellino et al., 2006***; ***Moreau et al., 2015***), because these signals would more efficiently shift cells between exploration scales.

## Supporting information

Figure1-VideoSupp1

Figure2-VideoSupp1

## Acknowledgments

We acknowledge Aya Ludin-Tal and Leonard Zon for the generous gift of the Tg(*lck*:GFP) zebrafish line. We acknowledge Kiran Kocherlakota and the Stanford VSC for assistance with zebrafish management and husbandry. We acknowledge Saroja Korullu for assistance with library preparation. We also acknowledge Stanford Research Computing and the Sherlock2 computer cluster for computational support and resources. Finally, we acknowledge Louis Leung and Karen Mruk for invaluable advice regarding microscopy and zebrafish; Edward Marti for helpful discussions on imaging, analysis, and the manuscript, and Felix Hornes and Michael Swift for comments on the manuscript.

## Data and materials availability

Sequencing data and the gene expression count table have been deposited on GEO (accession: GSE137770). Analysis code and trajectory data are available at https://github.com/erjerison/TCellMigration.

## Materials and Methods

### Zebrafish lines and procedures

Tg(*lck*:GFP, *roy*^−/−^, *nacre*^−/−^) zebrafish (*Danio rerio*) (***Langenau et al., 2004***) were obtained as a generous gift from Dr. Leonard Zon and Dr. Aya Ludin-Tal. Imaging was performed on Tg(*lck*:GFP) zebrafish crossed into a *nacre*^−/−^ background, at between 9 and 13 dpf. All adult and larval zebrafish were maintained according to protocols approved by the Stanford Administrative Panel on Laboratory Animal Care.

### Microscopy

Imaging was performed on a single-plane illumination microscope constructed as specified in (***Pitrone et al., 2013***), with the exception that a Prior ProScan XY stage (Prior Scientific) coupled to a Zaber T-LLS 105 stage (Zaber Technologies) was used for sample movement. The light sheet was generated using an Olympus UMPLFLN10XW objective (NA=.3) and detection was performed with an Olympus UMPLFLN20XW objective (NA=.5) and an achromatic doublet tube lens (AC508-180-A-ML, Thorlabs). Images were recored either on a Retiga 2000R camera (Qimaging) or an Ace acA2040 (Basler). For the Ace ac2040 camera, a meniscus lens (LE1418-A-O2” N-BK7, Thorlabs) was added as a zoom lens, to match the image pixel width between the two cameras at .37 *μm*. The fluorescence source was an Obis LS 488 nm laser (Coherent), and the microscope was controlled by Micro-Manager.

Zebrafish between 9 and 13 dpf were anesthetized with Tricaine-S (MS-222, Pentaire; .008% w/v, buffered to pH 7) and embedded in 2% low melting point agarose (Lonza SeaPlaque, #50100) with .004% w/v Tricaine. For imaging, the agarose was submerged in E3 with .008% w/v Tricaine and 50 mM Hepes. With the exception of ***Figure 2*** and ***Figure 2***-***video 1***, tiled z-stacks were obtained every 45 seconds for at least 180 timepoints, with a field of view of at least 592 *μm* (dorsal-ventral axis) by 1200 *μm* (anterior-posterior axis). For ***Figure 2***A and ***Figure 2***-***video 1***, a z-stack was obtained every 12 seconds for 1100 timepoints, with a field of view of 757×568 *μm* (the first 900 timepoints are shown). For statistical comparison with the remainder of the data, trajectories from this final dataset were subsampled in time to give 48 second timesteps. Data was acquired with 2×2 binning, for an image pixel width of .74 *μm*.

For imaging in the presence of Rockout, embedded fish were submerged in E3 with .008% w/v Tricaine and 50 mM Hepes plus 12 *μm* Rockout (Sigma Aldrich #555553). For paired control/Rockout trials, fish were imaged for 2.5 hours in control conditions, followed by 2.5 hours in Rockout conditions over the same field of view.

### Single-cell RNA sequencing

Thirty 15 dpf Tg(*lck*:GFP) zebrafish were euthanized using .04% w/v Tricaine and transected posterior to the anus. Tail portions were pooled into HBSS (ThermoFisher #14025092) on ice. Tails were dissociated by incubating with 100 *μg*/*mL* Liberase-TL (Sigma Aldrich #5401020001) at room temperature for 20 minutes, followed by trituration with a 23 gauge needle. The cell suspension was filtered through a 40 *μm* filter and washed once in HBSS. GFP+ cells were sorted from an unbiased FSC-SSC gate on a Sony SH800 cell sorter into 384-well hard-shell PCR plates (Bio-Rad HSP3901) containing .4 *μl* of lysis buffer, prepared as described previously (***Schaum et al., 2018***). Reverse transcription following a Smart-Seq2 protocol, and Illumina library preparation, were carried out as described previously (***Schaum et al., 2018***), except that following cDNA amplification, cDNA was diluted uniformly to a mean target concentration of .4 ng_*μl* for library preparation. Libraries were sequenced on the NovaSeq 6000 Sequencing System (Illumina) using 2×100-bp paired-end reads.

### Image processing and cell tracking

Tiles were assembled based on recorded stage coordinates and a Maximum Z projection was applied to Z stacks. Sample drift in x and y was subtracted by identifying and tracking autofluorescent pigment spots. In particular, the coordinates of 1-3 isolated pigment spots were identified manually at the first timestep; at each timestep, the brightness centroid was computed for a circle with a 25 pixel radius around the previous centroid, and the average trajectory of the pixel spots was rounded to the nearest pixel and subtracted from the timeseries. Prior to cell segmentation, the average image across the whole timecourse was subtracted from each timestep. For data recorded on the Retiga 2000R camera, prior to segmentation the image was thresholded at the 30th pixel percentile and the maximum pixel value was fixed so that .4% of pixels were saturated. For data recorded on the Basler Ace acA2040 camera, no lower threshold was used and the maximum pixel value at each timepoint was fixed so that .2% of pixels were saturated. Ilastik software (***Sommer et al., 2011***) was used for cell segmentation and tracking: the Ilastik pixel classification module was used to classify foreground and background, and the manual tracking module was used to identify and track cells. To define trajectories, the brightness centroid of each cell in x and y at each timestep was computed from Ilastik tracking masks and the Maximum Z projection. Processing steps not using Ilastik were performed using Python 3.6 (code available at: https://github.com/erjerison/TCellMigration).

### Trajectory analysis

Trajectories with at least 30 consecutive steps were included in the analysis; for MSD calculations, trajectories that included all time intervals were included. For calculations of power spectra, single missing timesteps were linearly interpolated based on the two adjacent positions, and computations were performed on the longest consecutive segment for each trajectory. For the *M. musculum* data, the time interval was 30 seconds. For the *Dictyostelium* data, timesteps were subsampled from the original to give an interval of 20 seconds.

Mean-squared displacements were computed along each trajectory as:

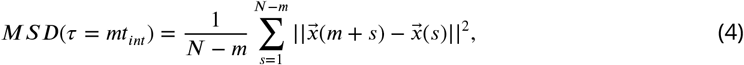

where *N* is the total number of timesteps and *t_int_* is the time interval. The overall MSD was computed by averaging the MSDs for each trajectory, and 95% confidence intervals were calculated via a bootstrap over trajectories.

The overall MSD was fit to:

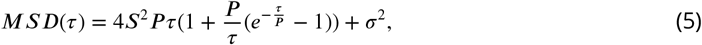

which we note is the common formula for mean squared displacement in both the OrnsteinUhelnbeck model (see Appendix 1) and in the Kratky-Porod wormlike chain model. Unless otherwise noted, fitting was performed using the scipy.optimize.curvefit function in scipy 1.3.0; fitting was performed in log space and weighted by computed confidence intervals.

The velocity power spectrum was computed based on the vector of secant-approximated velocities for each trajectory. Velocity vectors were zero-padded to 400 timesteps, and the fourier transforms of the velocity components were computed using the the fft function in numpy (1.16.4). Letting the fourier-transformed velocity components for trajectory *m* be *v_x_*(*k*, *m*), *v*_*y*_(*k*, *m*), the power spectrum for each trajectory was computed as:

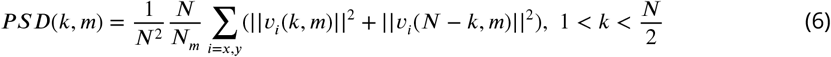

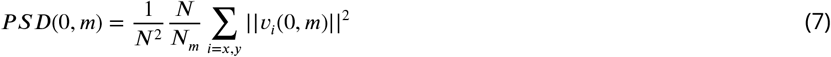

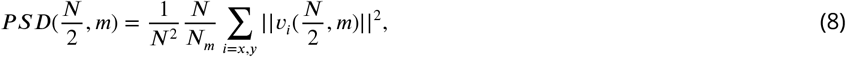

where *N* = 400 and *N_m_* is the length of trajectory *m*. The overall PSD was computed as the average over the PSDs for each trajectory:

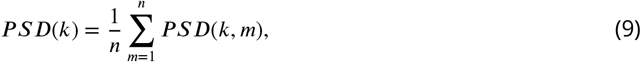

where *n* is the number of trajectories. For ***Figure 1***D, a piecewise linear function was fit to the PSD in log space; we plot the high-frequency fitted line and a line with slope 0.

Following (***Petrovskii et al., 2011***), we calculated the distribution of bout lengths within a trajectory as the distribution of x displacements between reversals in direction in x, divided by the average of these displacements within each trajectory. The distribution was calculated using the numpy.histogram function on percentile bins with the option density=True; the x locations of points were determined based on the average value of points in each bin. We fit the distribution to a stretched exponential function 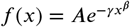; the fitted value of the stretch parameter *β* was .65.

The overall speed distribution was computed by collecting secant-approximated speeds across all trajectories and timepoints; similarly, the overall turn angle distribution was computed by collecting all relative angles between consecutive segments. For the Kolmogorov-Smirnov (KS) test, the overall CDFs of speeds and turn angles were estimated by measuring the cumulative frequency over 25 percentile bins and performing linear interpolation to yield a continuous function. A two-sided KS test (scipy.stats.kstest) was performed for the sets of speeds and turn angles of each trajectory.

Turn angle distributions for each speed class were computed by collecting all relative angles between consecutive segments amongst cells in that speed class; the distributions were symmetric about *θ* = 0 and so were folded to be between 0 and *π* radians. 95% confidence intervals were calculated based on a bootstrap over trajectories in each speed class. For the relationship between local speed and turn angles (***Figure 2***B), the local speed was estimated as the average speed of the two consecutive steps surrounding a turn. Turns were binned based on the local speed, and the average of the cosine of the turn angles was computed for each bin. For this and other binned statistics, the x location of the bin was fixed to be the average value for the points in that bin.

To estimate the rate of speed switching, all trajectories of at least 120 minutes in length were used. The average speed of each trajectory was measured on 20 minute intervals; 20 minutes was chosen to minimize the bias-variance trade-off. Specifically, because every cell samples speeds from a distribution, there is trade-off between measuring speeds on intervals that are too short, which may not give a good estimate of the mean, and intervals that are too long, where cells may switch during the interval. To minimize this trade-off, the interval that maximized the rank correlation between adjacent non-overlapping blocks was used. The average speed of each cell was measured on non-overlapping intervals, and the Spearman rank correlation coefficient between all pairs of intervals was computed. The correlation as a function of time was calculated as the average over all pairs of intervals with the same difference in start times. We computed 95% confidence on a bootstrap over trajectories. For the null model, we permuted speeds amongst the trajectories on each interval; we calculated 95% confidence intervals over the permutations.

For ***Figure 3***, the average of the cosine of turn angles between adjacent steps was calculated for each trajectory, as well as the average over all adjacent steps of the secant approximated speeds. For ***Figure 3***D, the correlation between local speed and turns was computed as the Pearson correlation coefficient between the local speed, as defined above, and turn angles across the set of adjacent steps in the trajectory.

To estimate the fraction of the variance in turning behavior explained by the cell speed, we fita spline curve (UnivariateSpline class of scipy 1.3.0; default parameters) to the relationship between speed and the average of the cosine of the turn angles (***Figure 3***-***Figure Supplement 1***A). Letting the spline function be *f*, we estimated the variance accounted for by the speed as *V_s_ = V ar(f(S_m_))*, where the index *m* labels trajectories. We estimated the variance in the means due to variation within trajectories, which we called stochasticity, as 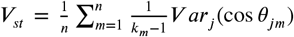, where *n* is the total number of trajectories, *k*_*m*_ is the number of turn angles within trajectory *m*, and cos *θ*_*jm*_ is the cosine of the turn angle *j* in trajectory *m*. Remaining variance we classified as other (***Figure 3***-***Figure Supplement 1***B); this may be due to imperfections in the spline model, other experimental noise, or additional biological variability.

The persistence time was defined to be the time elapsed before the trajectory turns at least 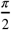 radians, averaged along the trajectory. Specifically, letting the displacement between timepoints *s* and *s* +1 be 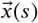, the persistence time along each trajectory was calculated as:

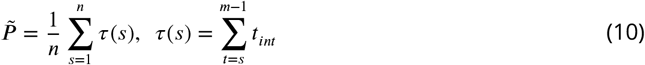

where *m* > *s* is the first timestep for which 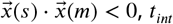 is the time interval, and *n* is the final base point for which *m* ≤ *N*, where *N* is the final timepoint. For ***Figure 4***A, trajectories were binned into mean speed deciles, and the average persistence time was calculated over trajectories in the bin; error bars represent 95% confidence intervals on a bootstrap over trajectories.

The effective diffusion coefficient at time *τ* was measured as:

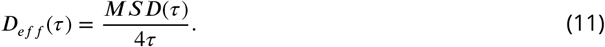

To measure *D*_*eff*_(*τ*) as a function of *S*, cells were divided into speed bins (with 5% of the speed distribution per bin); *D_eff_*(*τ*) for each speed bin was measured by averaging the *D_eff_*(*τ*) across trajectories, and error bars were computed based on a bootstrap over all trajectories. Note that *D_eff_*(*τ*) will be independent of *τ* only if diffusion scaling is respected, so that the collapse of the data in ***Figure 2***F is additional corroboration that the trajectories behave diffusively at long times.

As shown in the models section of the SI below, under the UPT model:

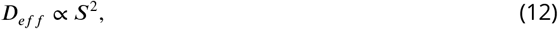

whereas under the SPC model,

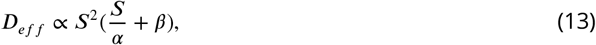

where *α* and *β* are fixed across all trajectories. We fixed *α* and *β* by fitting a line to the persistence time relationship in ***Figure 1***C; note that this is a short-time statistic and need not a priori predict the effective diffusion constant at longer times. We fit the UPT model (dashed line) and the SPC model (solid) to the measured *D_eff_* as a function of S; in both cases, there was one fitting parameter which was the constant of proportionality, which allows for an offset on the y-axis in log space but does not change the shape of the curve.

### Analysis of scRNAseq data

Reads were aligned to the Zebrafish reference genome (genome release: GRCz10; annotations: GRCz10.85) using STAR (2.5); reads aligned to each gene were counted using the htseq-count function of HTseq (0.8.0), with the options-m intersection-nonempty and -nonunique all. Note that the final option counts reads that align to a location with more than one annotated feature (e.g. overlapping ORFs) as belonging to both features. This is necessary because of mis-annotation of the T cell receptor light chain constant region in the Zebrafish reference genome; both ENSDARG00000075807 (*traj39*) and ENSDARG00000104132 (*traj28*) contain the *trac*, so that reads mapping to *trac* would otherwise be discarded.

Cells were filtered if they expressed fewer than 650 genes or more than 3250 genes, and if more than 8% of reads were of mitochondrial origin. We used UMAP (0.3.1) with the default options to embed the log-transformed counts table in two dimensions, and HDBSCAN (0.8.22) with min_samples=10 to call clusters (Figure5-***Figure Supplement 1***A). Comparison with the index sort data (Figure5-***Figure Supplement 1***B) showed that cells from the largest cluster had FSC-BSC consistent with lymphocytes, whereas cells from other clusters tended to have higher FSC and BSC. We called the major cluster as the first cell group and other clusters as the second group; the n=23 cells that were not assigned to a cluster by HDBSCAN were included with the first group if they had BSC < 8 × 10^4^, and with the second group otherwise. We identified differentially expressed genes between the two groups via a Wilcoxon Rank-Sum test. We selected the 50 most differentially-expressed genes (lowest Wilcoxon p-value) that were enriched in the T cell group for further analysis. These included the T cell and immune-related genes *tagapa*, *tagapb*, *ccr9a*, *tnfrs9b*, *il2rb*, and *ptprc* (***Schaum et al., 2018***); we also tested for expression of the T cell receptor light chain constant region (*trac*; expression estimated based on the mean expression of ENSDARG00000075807 and ENSDARG00000104132). Based on these markers, we identified the larger group (n=351) as T cells, and show expression for these cells in ***Figure 5***B. The 50 most significantly differentially expressed genes enriched in this group also included *arpc1b*, *wasb*, *arghdig*, *coro1a*, *sept9b*, and *capgb*, which we classified as belonging to the WASP/ARP2/3 pathway based on the literature (***Vicente-Manzanares et al., 2002***). Finally, we observed very little expression of markers associated with other types of immune cells (the B cell light chain *igic1s*, the B cell marker *ccl35.2*, and the neutrophil and macrophage markers *mpeg1* and *mpx* (***Tang et al., 2017***) in either group (***Figure 5***B, ***Figure 5***-***Figure Supplement 2***). The genes most significantly enriched in the non-T cell group are primarily keratin proteins (*krt8*, *KRT1*), as well as the epithelial marker *ahnak* (***Schaum et al., 2018***) (***Figure 5***-***Figure Supplement 2***). We identified these cells as epithelial cells, likely keratinocytes. We note that we observed GFP signal in the somite region of the Tg(*lck*:GFP) tail via microscopy (see, e.g., ***Figure 1***A and Movie S1) which we did not observe in wildtype *nacre*^−/−^ zebrafish, suggesting that these cells may mis-express the marker.

Finally, we note that we observe a sub-cluster of T cells that are enriched for expression of *ccl38a.5*, *ccl38.6*, and *zbtb32*, and also have somewhat higher expression of *scinlb* (***Figure 5***-***Figure Supplement 2***). This is consistent with a subpopulation of cells previously identified in Tg(*lck*:GFP) zebrafish (***Tang et al., 2017***). These cells were previously identified as NK cells; however, since they express the T cell receptor light chain, they are likely to be a T cell subset.

## Appendix 1 Persistent random walks: the uniform persistence time (UPT) and the speed-persistence coupling (SPC) models

The infinitesimal model most commonly used to describe metazoan cell migration is the Orenstein-Uhlenbeck model (OU) (***Uhlenbeck and Ornstein, 1930***), which has also been called the persistent random walk model (PRW) in the context of cell migration (***Wu et al., 2014***). While we have chosen this model for concreteness, the statistical features discussed below are also in common to a number of other models that include some directional persistence but are diffusive at long times, including the Kratky-Porod wormlike chain model (***Doi and Edwards, 1988***). We briefly review some of the standard results which we use to compare to data below.

Under the OU model, the dynamics of a cell are described by:

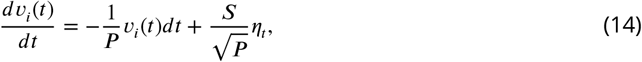

where S is the speed parameter; P is the persistence time parameter; *η*_*t*_ is a Gaussian white noise term; and *i* = *x*, *y*, *z*. This model, considered the prototypical noisy relaxation process, produces two main qualitative features: trajectories that turn smoothly (i.e. directional persistence), and diffusive behavior at times *t* ≫ *P*. We note that fluctuations in velocity and speed along the trajectory are also features of this model. In particular, the velocity-velocity autocorrelation function is given by:

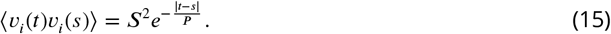

Setting *t* = *s*, we see that the speed parameter *S* is proportional to the root-mean squared speed: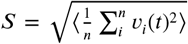, where *n* is the number of dimensions. Because the distribution of velocities generated by the model is Gaussian, *S* is also proportional to the mean speed. The decay of the velocity autocorrelation in each component sets the turning timescale at *P*; at long times the directions of motion are uncorrelated.

The mean-squared displacement (MSD) after a time interval *τ* is given by:

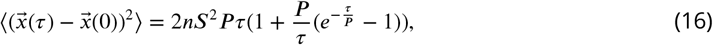

where n is the number of dimensions. The MSD scales advectively, as *nS*^2^*τ*^2^, in the limit of *τ* ≪ *P*, and diffusively, as *2nS*^2^*Pτ*, in the limit of *τ* ≫ P. Thus the model predicts that 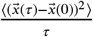 will approach a constant value of *2nS*^2^*P* at long times, which defines the effective diffusion constant to be 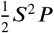.

We refer to the OU model with fixed persistence time parameter *P*(but potentially variable speed parameters *S*) as the uniform persistence time (UPT) model. We note that under the OU model, the MSD and PSD depend on the speed parameter *S* only through the constant scale factor *S*^2^: for fixed *P*, the quantities 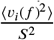 and 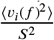 are independent of speed, as are the normalized quantities 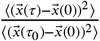 and 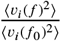, where *τ*_0_ and *f*_0_ are a chosen time interval and frequency, respectively. that this is also(tNruoteeof the full dynamics: we can eliminate the dependence on *S* by transforming to the variable 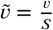, or measuring distance in units proportional to *S*.) In particular, the effective diffusion constant 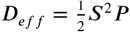 scales with *S*^2^.

Our observation of a linear relationship between measured mean speeds and correlation times suggests the following constrained form of the OU model, which we have called the seed-persistence coupling (SPC) model:

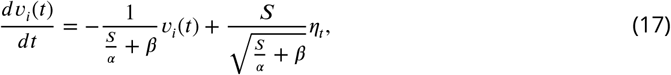

where *α* is a constant with units of acceleration; *β* is a constant with units of time; and both are fixed across all cells.

In this model, the effective diffusion constant is:

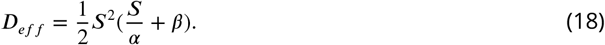

Under the SPC model, the control parameter *S* is proportional to the cell’s mean speed, so that this observable fully specifies its dynamics.

Finally, we note that with fixed *S* and *P*, these models still produce variation in both local speed and turning behavior along trajectories; the control parameters, together with Equation 14, set the distributions of these quantities.

## Efects of finite-length trajectories, sampling intervals, noise, and distributions of persistence time on speed-persistence coupling

Measurements of speed and persistence time are imperfect estimators of the underlying continuous process. Here we address whether statistical artifacts could generate the observed correlations between speed and persistence time. In particular, the finite sampling interval introduces a bias downwards in all speed estimates, because some turns are missed. Because this effect is stronger for less-persistent cells, which turn more, we expect it to introduce a correlation between the measured speed and the measured persistence time.

To evaluate the influence that this may have had on our data, we simulated a collection of cells with the same speeds as our measured cells under the OU model. Simulations were performed using the velocity update rule in Equation 14 (***Gillespie, 1996***), with 20 simulated intervals dt per sampling interval. Position coordinates were determined by numerical integration of the velocities along the simulated trajectories. Noise in centroid locations was included by adding a Gaussian random variable to x and y positions. We conservatively set the noise parameter at *σ* = 3 *μm* per sampling interval; this was chosen as an estimate of the combined effects of true technical noise and changes in cell shape during the interval. Each simulated trajectory was 50 sampling intervals (1000 microscopic timesteps) in length. To match the measurement, sampling intervals were assigned to be 45 seconds in length. We measured both mean speeds and correlation times on the simulated trajectories as defined in Materials and Methods.

We first assessed whether the SPC model, defined in the previous section, with the addition of noise in the centroid locations, gave the expected dependence of measured persistence time on measured cell speed (Figure 7B). Next, we simulated a collection of cells with the same set of speeds as in the data, with a constant persistence time parameter P (the UPT model), to check whether the finite length of the trajectories induced a correlation between measured speeds and persistence times (Figure 7C). We did not observe a significant effect. Next, we added centroid location noise to the UPT model. The addition of noise does induce a correlation at the slow end of the speed spectrum (Figure 7D); this is because cells that happen to have turned more will appear slower. However, this effect becomes negligible for cells with speeds above the noise level. We note that this likely contributes to the measured propensity for sharp turns amongst the slowest cells, and may lead to misassignment between the two lowest speed classes.

**Figure 7.**
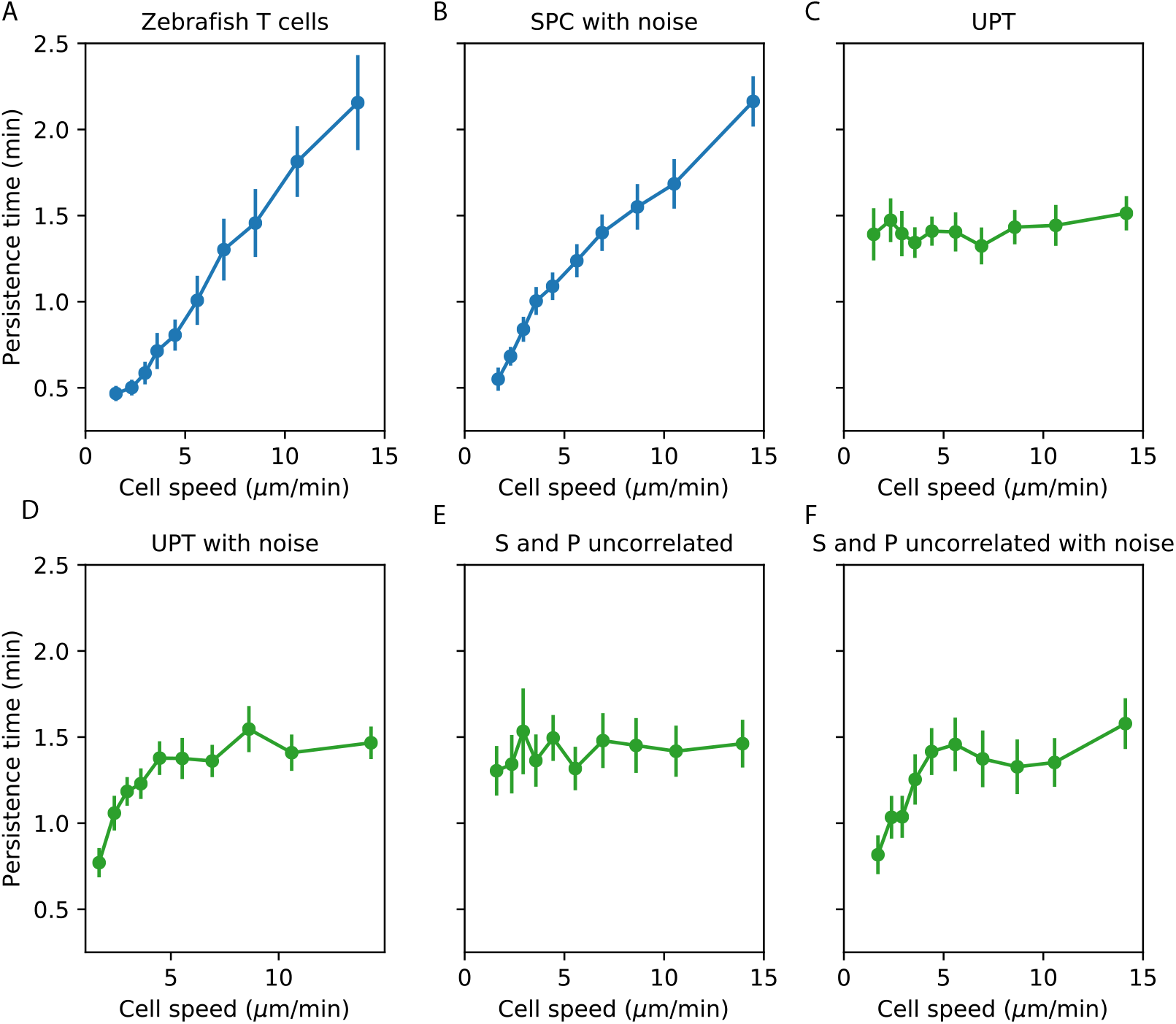
Comparisons between speed-persistence time relationship in simulations and data. A. Data (as in ***Figure 4***A). B. Simulation of the SPC model with empirical parameters (see Appendix 1 for details). C. Simulation of uniform persistence time (UPT) model, with speeds and persistence times measured as in the data. Biases introduced by measured speeds and persistence times do not lead to an observable correlation. D. Simulation of UPT model with a conservative estimate of mislocation noise. A spurious correllation is induced at low cell speeds, but cannot account for the trend across cell speeds that we observe. E. Simulation of a model where the predicted persistence times have been reshuffled amongst the cells, to simulate an empirically-realistic model where persistence times vary but are uncorrelated with speed. This does not generate a significant bias in the speed-persistence relationship. F. Model with reshuffled persistence times, as in E., and mislocation noise. As in D., this leads to a spurious correlation at low speeds but no other significant effects.

We next evaluated whether a model where both *S* and *P* varied in a manner consistent with the data, but were uncorrelated with each other, could induce a correlation between measured speed and persistence time. Such a correlation could appear on the faster end of the speed spectrum due to variable *P*, because the fastest measured cells are biased to having been both fast and particularly persistent. We evaluated the size of this effect in our data by simulating a collection of cells with the observed speed distribution, as before, permuting the predicted *P* parameters from the SPC model amongst the simulated cells. We found that this did not measurably bias the speed-persistence relationship (Figure 7E). Finally, we simulated the uncorrelated *S* and *P* model with the addition of centroid location noise (Figure 7F).

From this analysis, we concluded that noise in the locations creates a spurious correlation between measured speed and measured persistence time at the slow end of the speed spectrum, but that this effect cannot account for the consistent correlation across speeds that we observe; and that a model with variable *P* that is uncorrelated with *S* also cannot account for our observations.

**Figure 1–Figure supplement 1.**
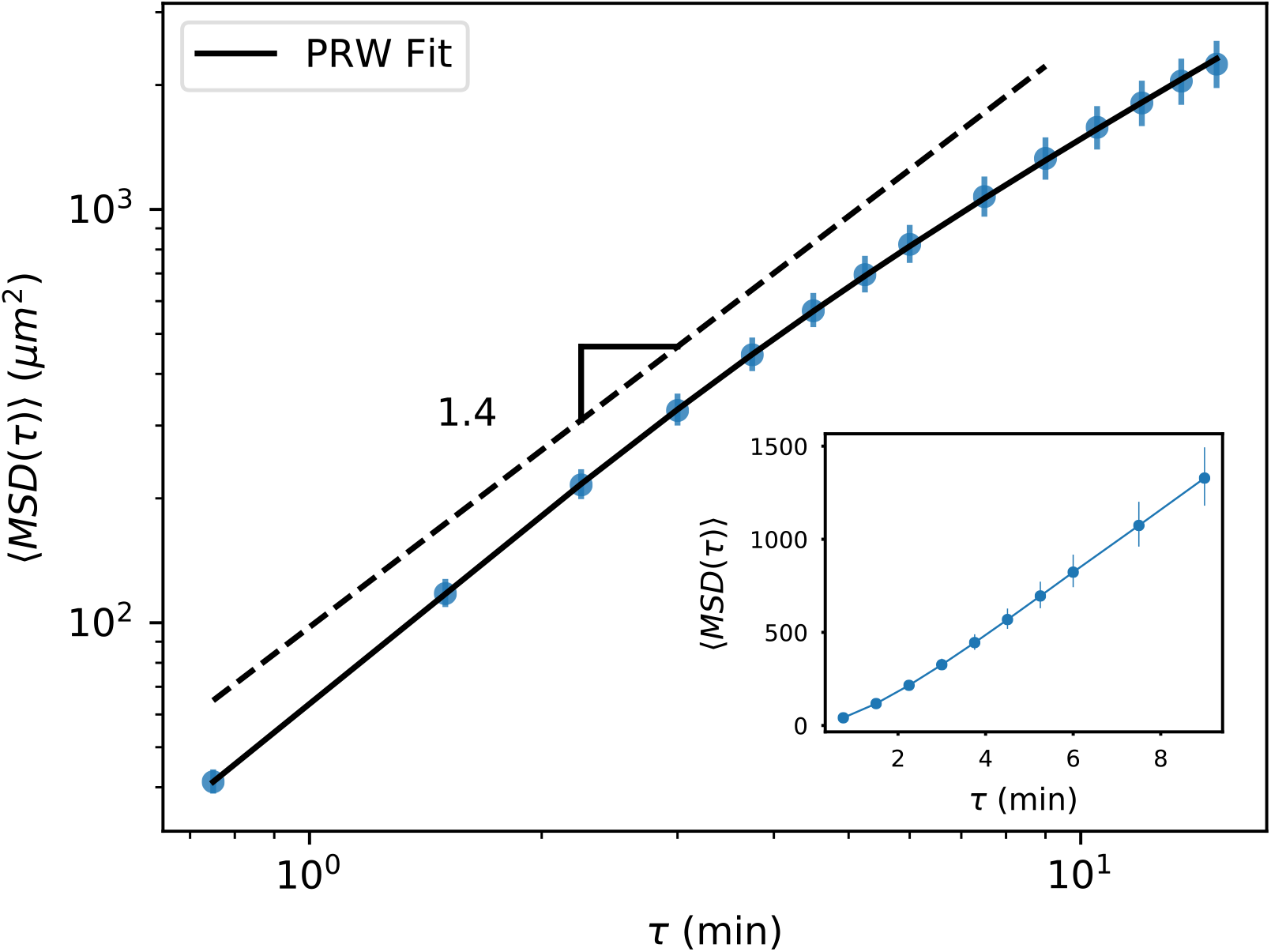
MSD for all trajectories tracked through 15 minutes, including all measured time intervals (n=612). As with the subset of longer trajectories, we observe a curved MSD consistent with a persistent random walk (PRW) model. Inset: linear scale through 10 minutes, showing a straight line consistent with diffusive behavior after the first few minutes. Error bars represent 95% confidence intervals on a bootstrap over trajectories.

**Figure 3–Figure supplement 1.**
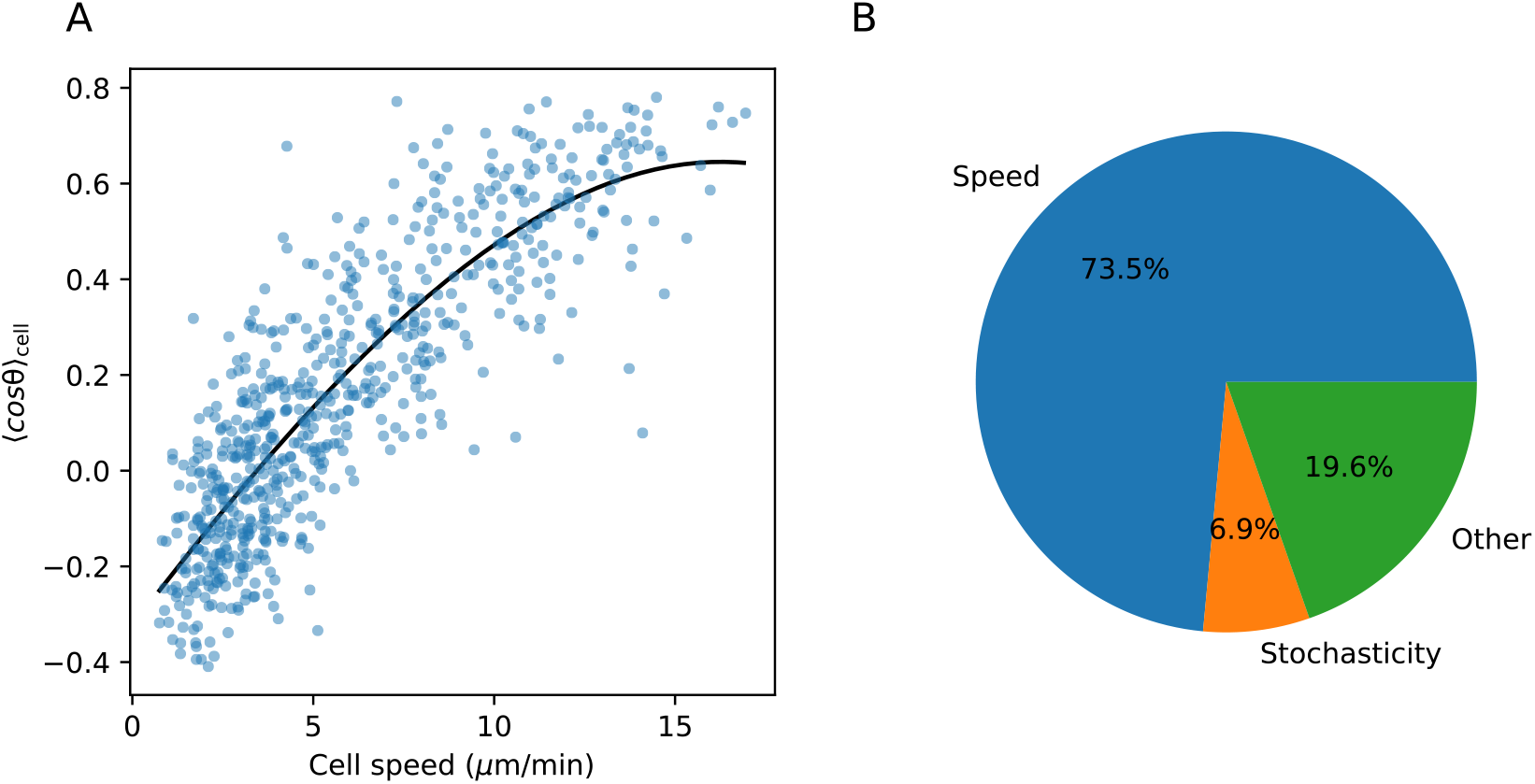
A. Spline fit to speed-turn angle relationship. B. The fraction of the variance in the turn angle summary statistic explained by speed (estimated based on the spline fit in A.), by stochasticity, i.e. variance within a trajectory; and by other factors, which may include imperfections in the spline fit (see Materials and Methods).

**Figure 4–Figure supplement 1.**
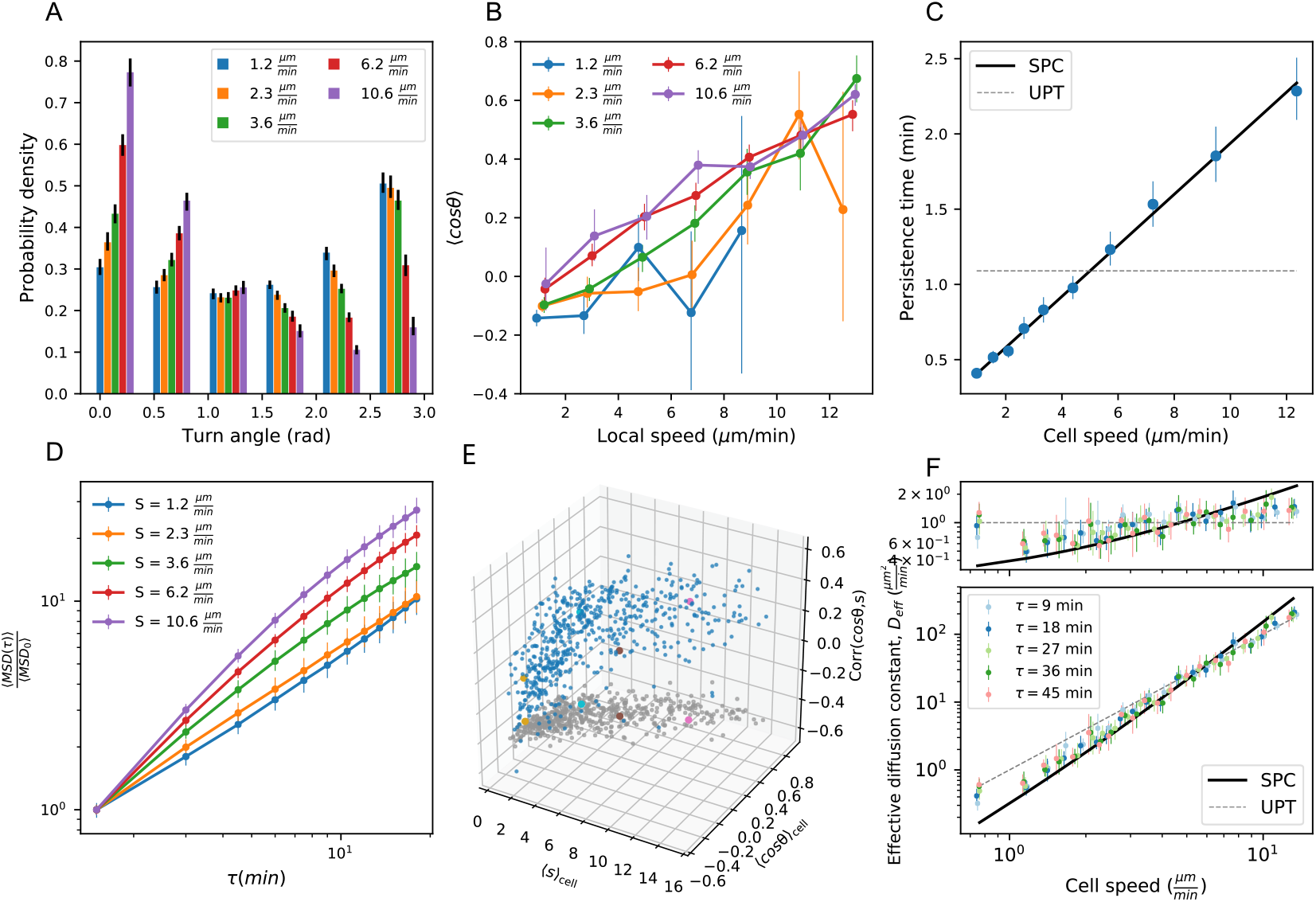
Panels A,B,D,E as in **Figure 3A-D**; Panels C,F as in **Figure 4A-B**; with all statistics re-calculated based on sub-sampling timepoints by a factor of 2.

**Figure 5–Figure supplement 1.**
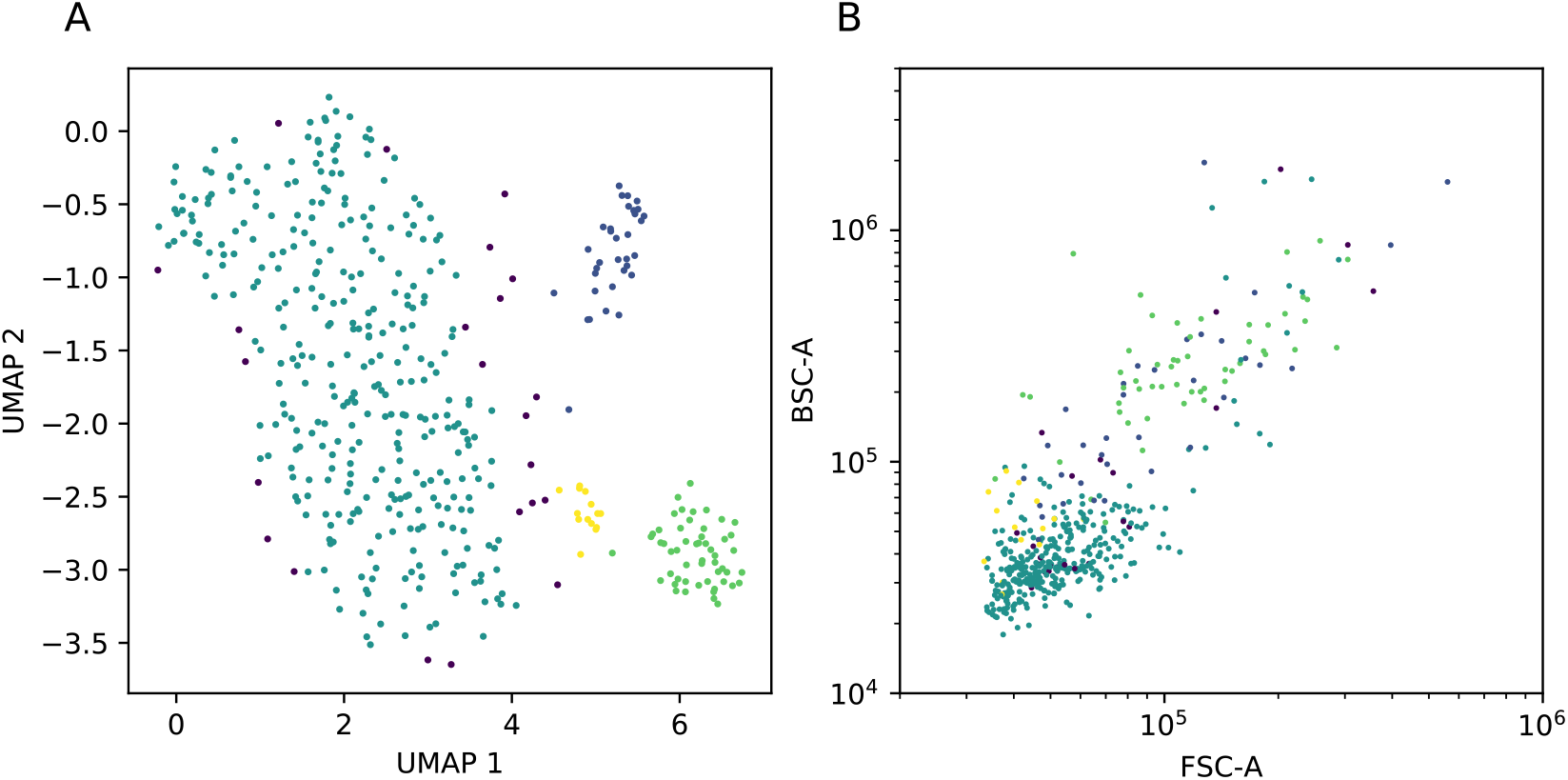
A. UMAP dimensional reduction of cell gene expression profiles from scRNAseq, with clusters assigned by HDBSCAN (colors). B. Index sort data of FSC/BSC for these cells (colors correspond to cluster assignments from A).

**Figure 5–Figure supplement 2.**
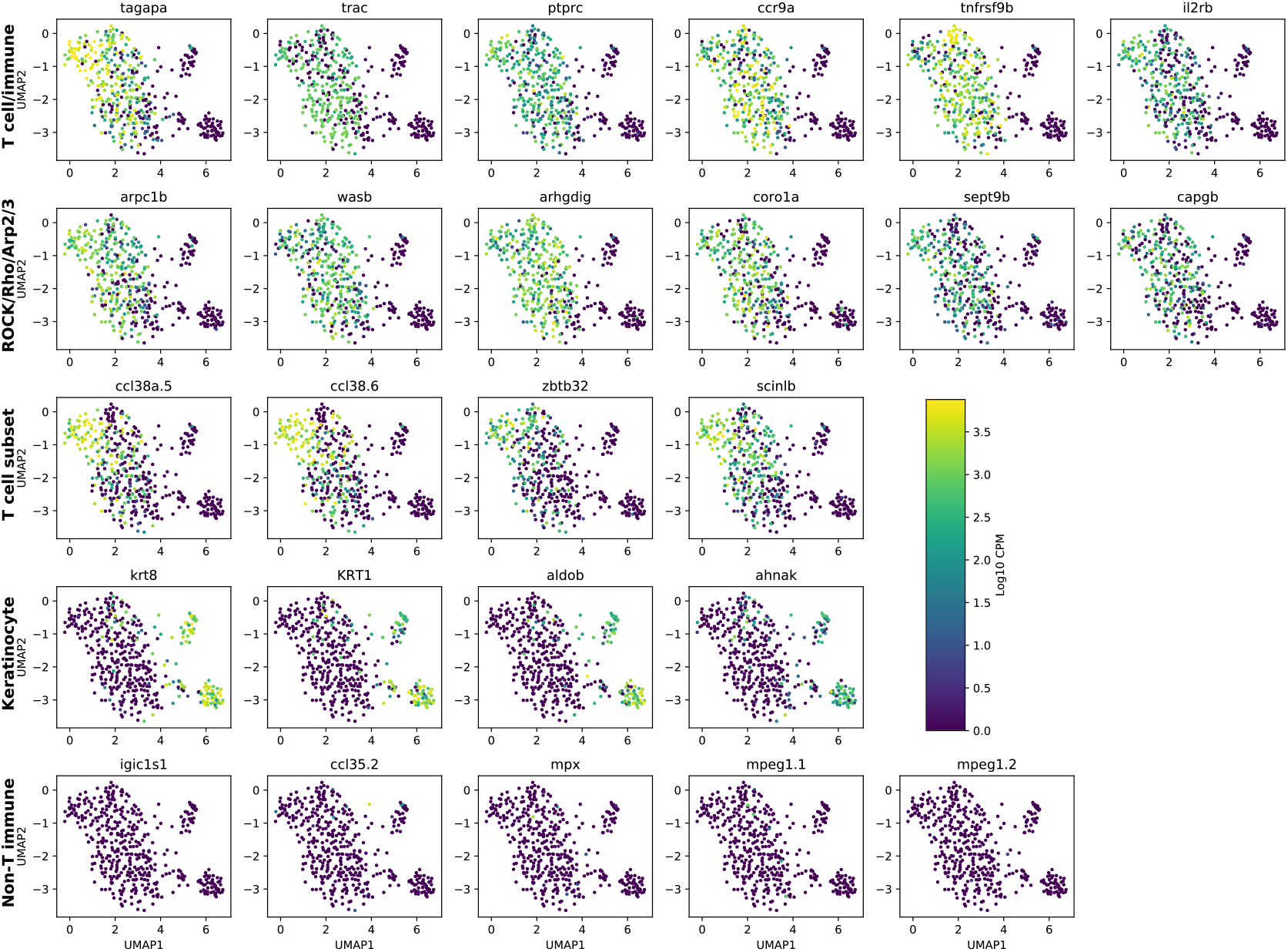
UMAP dimensional reduction as in Figure **Figure 5**, colored by expression of panels of genes. The largest cluster was called as T cells based on their expression of T cell and immune markers (first row) and non-expression of immune markers from other cell types (final row). The other cells were called as epithelial based on their expression of keratin genes, as well as *ahnak* (fourth row). We note that we observed a subset of T cells consistent with a subset from a previous observation (***Tang et al., 2017***) that were identified as NK cells (third row); however, these cells express *trac* as frequently as other T cells in the sample, suggesting that they are a T cell subset.

**Figure 5–Figure supplement 3.**
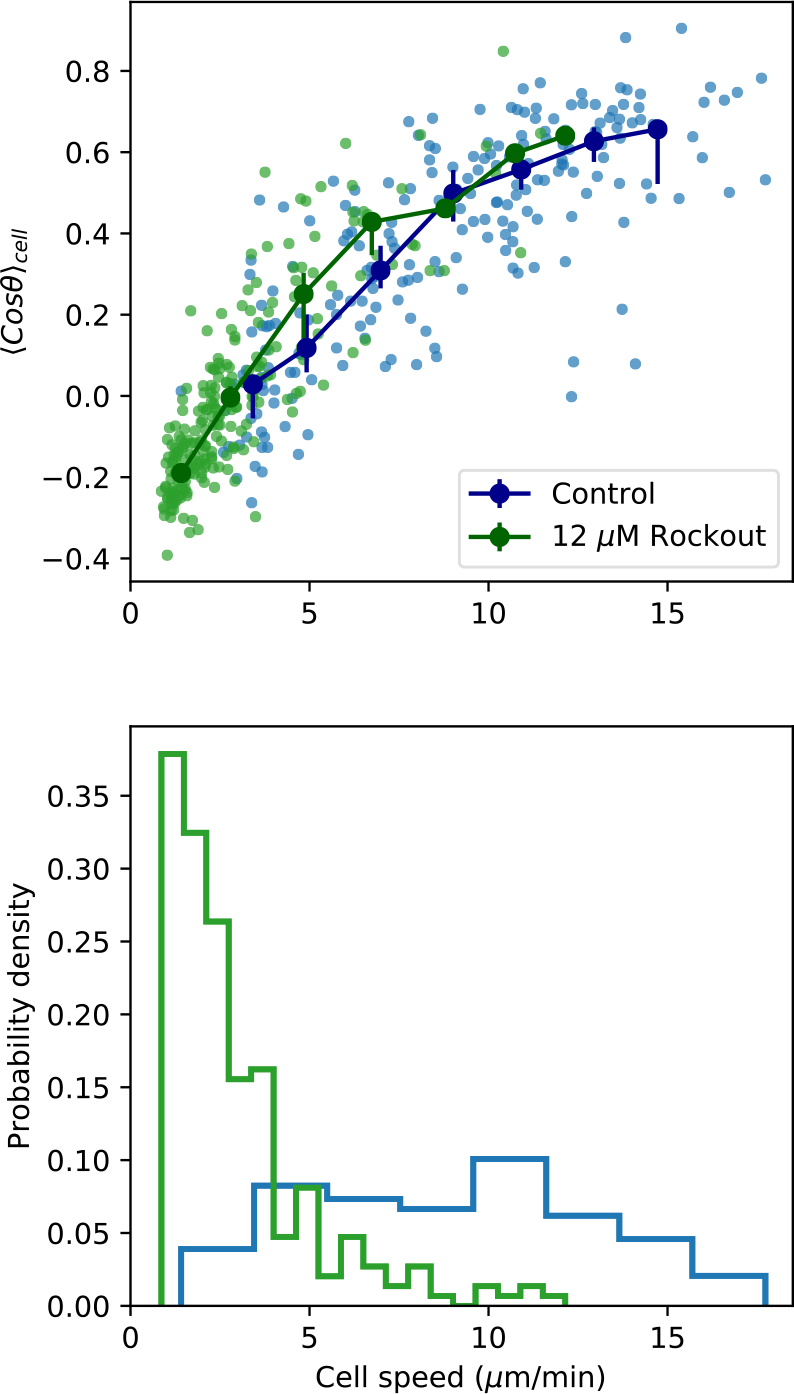
As in **Figure 5C-D**, but including only those control samples with a paired Rockout treatment sample (n=6 fish). Fish were imaged for 2.5 hours, and imaging media was replaced with media containing Rockout. Imaging over the same field of view was continued for 2.5 hours.

---

**Figure 1–Figure supplement 1**. MSD for all trajectories tracked through 15 minutes.

**Figure 1**–**video 1. T cell dynamics in the larval zebrafish tail and fin fold** Maximum Z projection of the tail of a Tg(*lck*:GFP, *nacre*^−/−^) at 12 dpf (GFP channel). Tiled Z stacks were recorded every 45 seconds for 2.5 hours (50 3 *μm* slices per stack). Tiles were assembled based on recorded stage locations. The movie was prepared using Python 3.6.0 (code available at: https://github.com/erjerison/TCellMigration).

**Figure 3–Figure supplement 1**. Variance explained by speed-turn relationship.

**Figure 4–Figure supplement 1**. Statistics from Figures 3 and 4, with timepoints subsampled.

**Figure 5–Figure supplement 1.** Comparison between UMAP and index sort.

**Figure 5–Figure supplement 2.**UMAP dimensional reduction as in ***Figure 5***, colored by expression of panels of genes.

**Figure 5–Figure supplement 3.***Figure 5 C-D*, including only paired control-Rockout treatment samples.

